# Primary cilia facilitate GPCR neurotransmission through the canonical cAMP cascade

**DOI:** 10.64898/2026.10.05.756873

**Authors:** Rita R. Fagan, Xinyi Jenny He, Nico Stuurman, Aliza T. Ehrlich, Mark von Zastrow

## Abstract

The primary cilium is increasingly recognized to be a discrete site for initiating neurotransmission through G protein-coupled receptors (GPCRs), yet how such neurotransmission is propagated downstream remains unclear. A canonical pathway of downstream GPCR neurotransmission is the cyclic AMP (cAMP) / cAMP-dependent protein kinase (PKA) cascade. While cAMP can be locally generated in cilia and produce important effects that are functionally compartmentalized there, neurotransmission through GPCRs must propagate to effectors outside of cilia. Thus, a prevailing current view is that ciliary neurotransmission is restricted to non-canonical pathways, or to cAMP effects confined within the cilium. Revisiting this view, we show here that the primary cilium of striatal medium spiny neurons is a major site from which endogenous D1 dopamine receptors initiate cAMP and PKA activity elevations that propagate throughout the extraciliary cytoplasm. We also show that the primary cilium spatially aligns essentially this entire ‘long-range’ signaling axis– from intercalating with numerous dopaminergic axon varicosities that initiate neurotransmission to enforcing proximity with somatic PKA stores that are activated by cAMP downstream. Together, these results extend the concept of ciliary neurotransmission to dopaminergic signaling in striatal neurons and reveal that, in this system, the primary cilium facilitates– rather than restricts– neurotransmission through the canonical cAMP / PKA cascade.

## Introduction

Nearly all cells in the mammalian nervous system project a solitary primary cilium^1^. Cilia have well-known signaling functions in the nervous system, most notably in enabling efficient phototransduction through cyclic nucleotides in specialized sensory neurons^2–4^ and supporting developmental signaling through the Hedgehog pathway in neural progenitor cells^5–7^. However, many other cell types in the nervous system, in addition to sensory neurons and neural progenitors, project a cilium and maintain it throughout life, thereby raising the fundamental question of whether neuronal primary cilia have additional functions^8^. The discoveries that neuronal primary cilia contain key GPCR signaling proteins such as specific adenylyl cyclases^9,10^, and display on their surface a select subset of GPCRs that are activated by classical modulatory neurotransmitters– including somatostatin^11^, serotonin^12,13^ and dopamine^14,15^– motivated the hypothesis that primary cilia represent a discrete site on the neuronal cell body (soma) for initiating modulatory neurotransmission through GPCRs^16^.

This hypothesis has gained support from *in vivo* studies demonstrating that genetic knockout or inhibition of signaling through cilia-localizing GPCRs^17,18^, or disrupting the primary cilium itself^19,20^, produces physiological and behavioral effects consistent with neuromodulatory dysfunction. Additional support has come recently through studies using sophisticated volumetric imaging methods, which have revealed that neuronal cilia (as well as cilia projecting from some other cell types) are frequently located close to conventional anatomical synapses and neurotransmitter-containing axon varicosities. In addition, it has been shown using *in situ* biosensors that GPCRs localized on the neuronal primary cilium can sense not only the ambient concentration of modulatory neurotransmitter, but also the dynamic release of transmitter from local axon varicosities^21,22^.

A key question moving forward is how such neurotransmission is propagated downstream from the cilium. A canonical pathway of downstream neurotransmission through GPCRs involves receptor-triggered production of cyclic AMP (cAMP) that acts as a diffusible mediator to drive activation of the cAMP-dependent protein kinase (PKA) that, in turn, controls a broad network of effector proteins which are distributed throughout neurons at multiple cytoplasmic and membrane sites^23–25^. While primary cilia are well known to contain adenylyl cyclases and support local cAMP production, many signaling effects of ciliary cAMP production are physically or functionally compartmentalized within the primary cilium rather than being broadly propagated^26–29^. Moreover, some previous studies have reported that GPCRs which classically initiate neurotransmission through the cAMP / PKA cascade, such as the D1-type dopamine receptor (DRD1 or D1R)^30,31^, have little signaling activity in the primary cilium^21,32^. Accordingly, a prevailing current view is that downstream neurotransmission initiated by GPCR activation in the cilium is necessarily restricted to noncanonical, cAMP-independent pathways^33–35^.

Here we revisit this view in light of prior evidence that the D1R can indeed stimulate ciliary cAMP production when over-expressed in a non-neural cell model^36^. We focus on striatal medium spiny neurons (MSNs) because many of these neurons natively express D1Rs, which have been shown to be selectively targeted to the primary cilium of MSNs both in cultured neurons and in vivo^14,15^, and D1Rs mediate neurotransmission in MSNs largely through the canonical cAMP / PKA cascade^23^. By devising a method to inhibit receptor activity locally and selectively in the ciliary microdomain, our studies reveal that the primary cilium is a major site from which endogenous D1Rs initiate the elevation of cAMP and PKA activity throughout the soma. We conclude that, in striatal MSNs, the primary cilium facilitates– rather than restricts– endogenous GPCR transmission through the canonical cAMP / PKA cascade and we propose a simple model for how this is achieved.

## Results

### Primary cilia on MSNs are intricately intercalated with dopaminergic axon varicosities *in situ*

In adult mouse striatal sections we localized neuronal primary cilia, marked by adenylyl cyclase type 3 (ADCY3), in close proximity with numerous dopaminergic axon varicosities marked by dopamine transporter (DAT, **Fig 1A**). Each primary cilium appeared to be embedded in a dense meshwork of dopaminergic axon varicosities (**Fig 1A’**) such that, on average, each cilium localized within 1 µm of seven resolved varicosities (**Fig 1A’’** and Supplemental Fig 1A-B). This is consistent with the facts that neuronal primary cilia in the striatum are unusually long (∼10 µm^37,38^) and that dopamine-containing axon varicosities are densely distributed^39^, but we also note recent evidence for specific tropism of dopaminergic axons to D1R-expressing MSNs^40^. Further, when one considers the rapid diffusion rate of dopamine in striatum (>400 µm^2^ s-1^39^), the number of potential release sites sampled by each cilium is likely higher than this estimate. Thus, each cilium is poised to detect dopamine released from numerous sites, provided that the relevant receptors are there.

**Figure 1.**
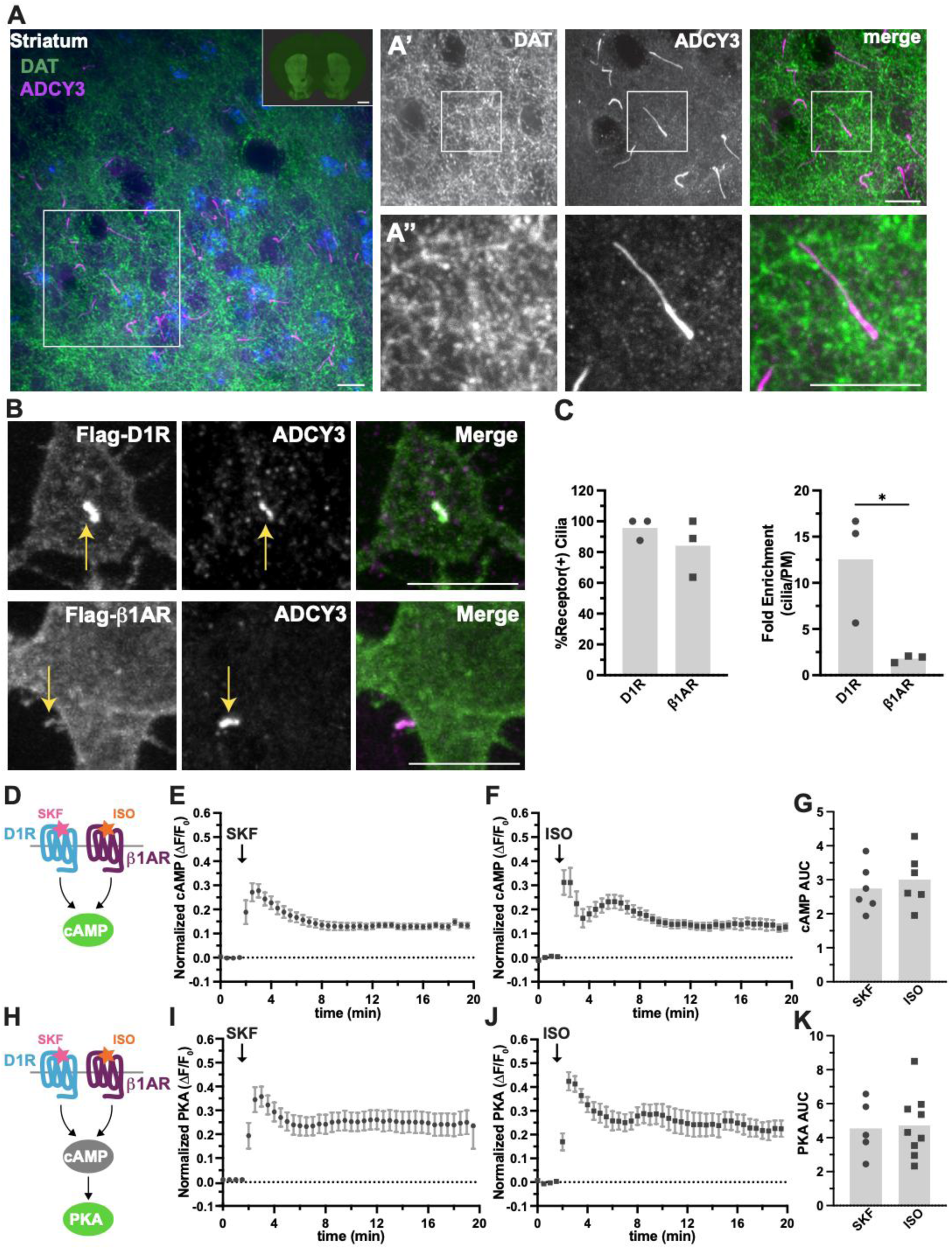
Primary cilia closely appose dopamine terminals and enrich D1R in striatal MSNs. **A.** Representative confocal image of neuronal primary cilia (ADCY3), dopamine terminals (DAT), and nuclei (DAPI) in adult mouse dorsal striatum. Scale bars: 100 µm. Inset: widefield image of DA terminals (DAT) of a whole 25µm coronal slice. Scale bar: 1000 µm. **A’.** Higher magnification and single channel views of the field highlighted in A. Scale bar: 10 µm. **A”.** Further magnification of the fields highlighted in A’. Scale bar: 10 µm. **B.** Representative confocal images of MSNs expressing Flag-D1R (top) or Flag-ꞵ1AR (bottom). Cells were incubated with anti-M1 (FLAG) antibody to label surface receptors prior to fixation and permeabilization and cilia were labeled with anti-ADCY3 antibody. Arrows indicate cilia-localized receptors. Scale bars: 10 µm. **C.** The percent cilia-localized receptor was similar for D1R and ꞵ1AR (left). D1R was significantly more enriched within cilia than ꞵ1AR (right). Data are presented as mean of 3 independent experiments, 26-28 cells, Unpaired t-test * p < 0.05. **D-G.** Cultured MSNs expressing cADDis biosensor were stimulated with 1 µM SKF (**E**) or 100 nM ISO (**F**), and fluorescence was measured over time by confocal microscopy. Background fluorescence is subtracted and data are normalized to maximal sensor response as defined by FSK+IBMX treatment after 20 min. **G.** ISO and SKF stimulated cAMP to similar degrees in striatal MSNs as measured by area under the curve (AUC); data are presented as mean ± SEM from 6 dishes from 3-4 independent neuron preps, 76-122 cells. **H-K.** Cultured MSNs expressing ExRai-AKAR2 were stimulated with 1 µM SKF (**I**) or 100 nM ISO (**J**), and fluorescence was measured over time by confocal microscopy. Data are normalized to maximal sensor response as defined by FSK+IBMX treatment after 20 min. **K.** SKF and ISO stimulated PKA in neurons to similar degrees as measured by AUC. Data are analyzed using Unpaired t-test and presented as mean ± SEM from 5-9 dishes from 3-5 independent neuron preps, 61-84 cells.

### Both D1 dopamine and β1-adrenergic receptors accumulate on primary cilia, but do so to different degrees

Previous studies have demonstrated selective targeting of D1R to the primary cilium, both in transfected non-neural cells and in striatal neurons^14,41^, and selective targeting of endogenous D1R to the primary cilium of MSNs has been demonstrated in the intact striatum^15^. We verified localization of a Flag epitope-tagged D1R to the primary cilium when expressed in cultured MSNs, and also verified its visible concentration on the cilium relative to the adjacent extraciliary plasma membrane where it was also detected (**Fig 1B, C**). We next tested the β1-adrenergic receptor (β1AR), a closely related GPCR paralog that is also endogenously expressed in MSNs and initiates neuromodulation in these neurons through the cAMP / PKA cascade^42^. Using the same experimental approach, we found a Flag-tagged β1AR to also localize on the primary cilium. However, unlike Flag-D1R, it was not detectably concentrated there relative to the extraciliary plasma membrane (**Fig 1B, C**).

As both D1R and β1AR mediate physiological neuromodulation in MSNs predominantly by activating the cAMP / PKA cascade^23,43^, we next verified native functional expression of each GPCR by assaying the regulation of cytoplasmic cAMP levels using cADDis, a real-time fluorescent cAMP biosensor that localizes diffusely throughout the soma in MSNs^10^. We used SKF81297 (SKF) to selectively activate D1R and isoproterenol (ISO) to activate β1AR (**Fig 1D**), choosing concentrations that produce near-maximal cAMP responses selectively through each (1 and 0.1 μM, respectively). SKF-induced activation of D1R produced a rapid cAMP elevation followed by a decay to a plateau level (**Fig 1E**), consistent with previous studies^10^. ISO-induced activation of β1AR elicited a qualitatively similar response (**Fig 1F**). To compare the magnitude of D1R and β1AR-mediated cAMP elevations across recordings and experiments, we normalized each to that measured after applying the adenylyl cyclase activator forskolin (FSK) and phosphodiesterase inhibitor 3-isobutyl-1-methylxanthine (IBMX) to saturate the biosensor response^10^ (Supplemental Fig 1C-E). Using this strategy, we found the natively coexpressed D1R and β1AR drive the somatic cAMP elevation to a similar magnitude as well (**Fig 1G**).

We next assessed signaling further downstream, at the level of somatic PKA activity, using ExRai-AKAR2 as a well-established real-time biosensor^44^ (**Fig 1H**). Consistent with the cAMP results, both SKF (**Fig 1I**) and ISO (**Fig 1J**) rapidly elevated somatic PKA activity. When normalized to the maximal sensor response measured after application of FSK/IBMX, as described above, both responses were determined to be similar in magnitude (**Fig 1K** and Supplemental Fig 1F-H). These results indicate that both D1R and β1AR, despite differing significantly in their relative ability to concentrate on the primary cilium, produce a comparable activation of the canonical cAMP cascade when assessed at this key downstream step. Moreover, by targeting ExRai-AKAR2 specifically to the primary cilium, using fusion to Arl13b that is highly concentrated on the cytoplasmic leaflet of the ciliary membrane^45^ (cilia-ExRai-AKAR2), we found endogenous D1R and β1AR drive comparable PKA activity elevations when assessed locally within the primary cilium itself (Supplemental Fig 1I-L). These results are fully consistent with previous evidence su free diffusion of cyclic nucleotides^36,46–48^ and PKA catalytic subunits^49,50^ through the ciliary base, but led us to question the view that the primary cilium is restricted from contributing to modulatory neurotransmission through the canonical cAMP/ PKA cascade.

### Both cilia-localized D1R and β1AR pools contribute to the somatic cAMP elevation measured in MSNs

To further investigate this, we sought to selectively block signaling from the ciliary microdomain, focusing on the most upstream step in the pathway by directly blocking the signaling activity of the receptor. To do so, we devised a strategy using an engineered protein fragment derived from the Ras-like domain of Gs (mini-Gs or mGs), which is a signaling-inactive G protein mimetic^51^ shown previously to block Gs and Golf-coupled GPCR signaling when expressed at sufficiently levels in intact cells^52,53^. Rather than driving mGs to a high concentration globally in the cytoplasm, however, we employed a strategy to achieve selective concentration at defined membrane locations by fusing the Ras-like domain to a specific membrane targeting module^54^. We used an 11-residue sequence motif derived from the Lyn tyrosine kinase^55^ to target mGs to the plasma membrane overall, and Arl13b to drive its targeting specifically to the cilium (Lyn11-mGs and Arl13b-mGs, **Fig 2A**). By including mApple to enable the visualization of each construct by confocal microscopy, we verified in living neurons Lyn11-mGs localization along the entire somatic plasma membrane, both inside and outside of the cilium, and specific concentration of Arl13b-mGs in the cilium (**Fig 2B**). As a negative control for functional studies, we fused Lyn11 to mApple alone (without mGs) and verified its expected localization throughout the plasma membrane, both inside and outside of the ciliary microdomain (Supplemental Fig 2A, B).

**Figure 2.**
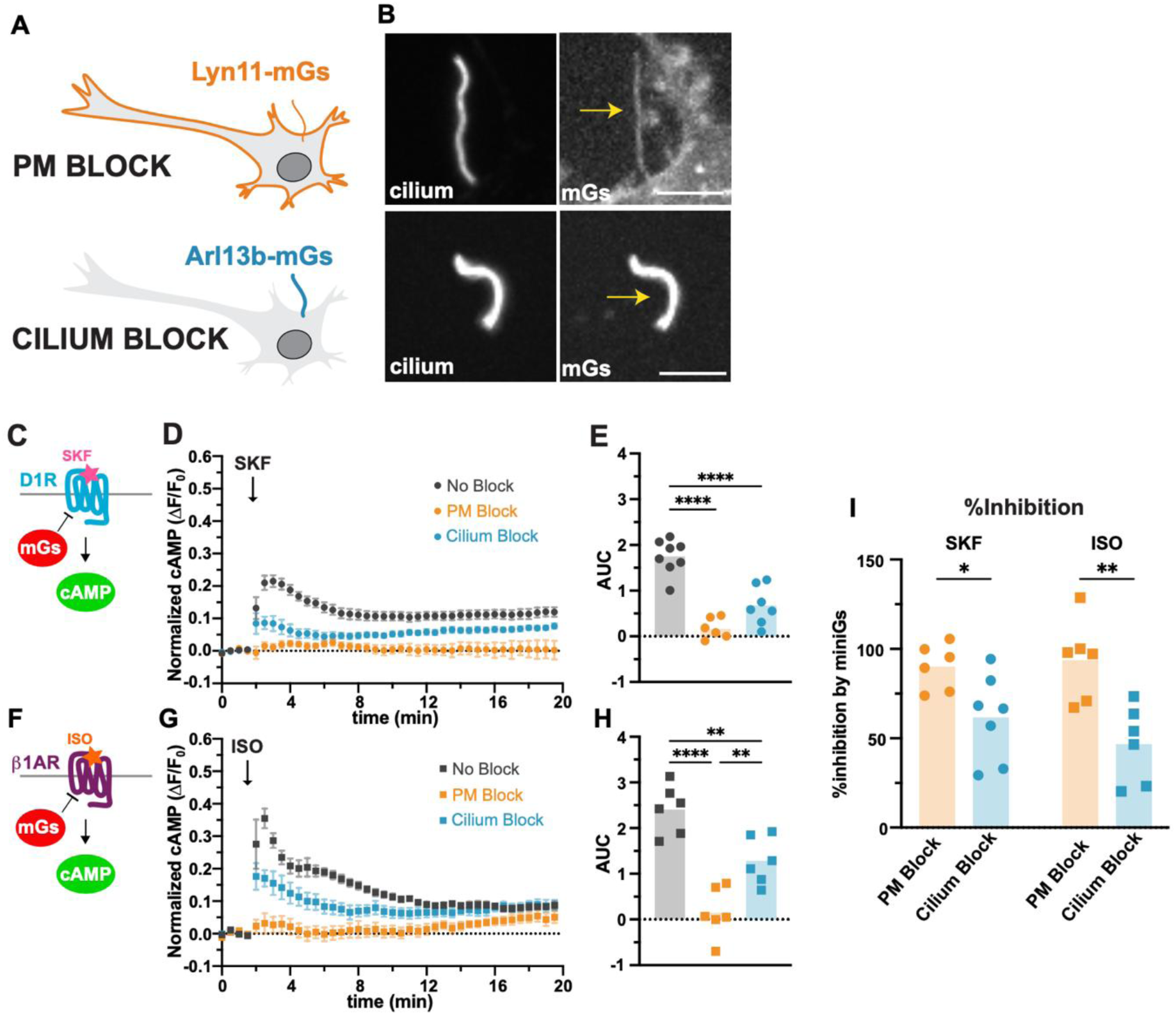
Targeted ciliary receptor blockade reveals a contribution of ciliary Gs signaling to dopamine and β-adrenergic receptor-evoked cAMP responses in striatal MSNs. **A.** Schematic describing the localization of the targeted mGs blocking constructs in neurons. **B.** Representative images of Lyn11-mGs (top) and Arl13b-mGs (bottom) co-expressed with Flag-HTR6 (left) to identify cilia. Lyn11-mGs is expressed along the plasma membrane and ciliary membrane, Arl13b-mGs is limited to the primary cilium. Arrows indicate mGs localized to primary cilia. Scale bars: 10 µm. **C-E.** Cultured MSNs expressing cAMP biosensor cADDis and Lyn11-mApple (‘No Block’), Lyn11-mGs (‘PM Block’), or Arl13b-mGs (‘Cilium Block’) were stimulated with 1 µM SKF and fluorescence was measured and quantified as described in Figure 1. **D.** Time course of cAMP response to 1 µM SKF added at 2 minutes in cultured MSNs expressing the different mGs blockers or control plasmid. **E. ‘**PM Block’ and ‘Cilium Block’ significantly reduced the integrated cAMP response to SKF (AUC) within the first 5 min following agonist addition in MSNs. One-way ANOVA ****p < 0.0001 with Tukey’s multiple comparison test; data are presented as mean from 6-8 dishes from 7 independent neuron preps, 65-170 cells. **F-H.** Cultured MSNs expressing control, ‘PM Block’, or ‘Cilium Block’ plasmid and cADDis cAMP sensor were stimulated with 100 nM ISO and fluorescence was measured and quantified as described in Figure 1. **G.** Time course of cAMP response to 100 nM ISO addition at 2 minutes in cultured MSNs expressing cADDis and ‘No Block’, ‘PM Block’, or ‘Cilium Block’ plasmids. **H.** Lyn11-mGs and Arl13b-mGs significantly reduced the integrated cAMP response to ISO in the first 5 min of agonist in MSNs. The ‘PM Block’ inhibited the cAMP response significantly more than the ‘Cilium Block’. One-way ANOVA ****p < 0,0001 with Tukey’s multiple comparison test; data are presented as mean from 6 dishes from 6-7 independent neuron preps, 65-103 cells. **I.** The degree of inhibition of the integrated cAMP response by ‘PM Block’ versus ‘Cilium Block’ was compared between SKF and ISO. The ‘Cilium Block’ inhibited cAMP significantly less than ‘PM Block’ for both SKF and ISO. Two-way ANOVA (interaction p = 0.28; main effect of mGs ***p = 0.0002; main effect of agonist p = 0.51) with Sidak’s multiple comparisons test. *p < 0.05, **p < 0.01, ***p < 0.001, ****p < 0.0001.

To assess the signaling effects of these blockers, we assessed the biosensor response integrated over the entire soma, bearing in mind that classical neurotransmission through the cAMP / PKA cascade requires signal propagation to effectors located outside of cilia^23^. We first assessed effects on the receptor-elicited cAMP elevation using the cADDis biosensor (**Fig 2C**). In neurons expressing Lyn11-mApple (the ‘No Block’ control condition), SKF produced a rapid elevation of cAMP (**Fig 2D**, black) similar to that observed in neurons not expressing it (compare with Fig 1E). Verifying the effectiveness of the inhibition strategy, the SKF-induced cytoplasmic cAMP elevation was almost completely suppressed in neurons expressing Lyn11-mGs (the ‘PM Block’ condition, **Fig 2D**, orange). In contrast, Arl13b-mGs (the ‘Cilium Block’ condition) only partially inhibited the measured cAMP elevation (**Fig 2D**, blue), producing a partial suppression when assessed by both the integrated AUC and peak metrics normalized to the FSK/IBMX maximum (**Fig 2E** and Supplemental Fig 2C).

We applied the same strategy to assess inhibition of the β1AR-mediated cAMP response (**Fig 2F**). Here again, the ‘PM Block’ almost completely suppressed the ISO-stimulated cAMP elevation but the ‘Cilium Block’ produced a partial effect (**Fig 2G, H** and Supplemental Fig 2D). We calculated % inhibition to compare the overall effects of the inhibitors between the receptors (**Fig 2I** and Supplemental Fig 2E). Together, these results suggest that both the endogenous D1R and β1AR are able to drive a detectable cAMP elevation in the soma from the primary cilium, but the cilium is not the only source because Arl13b-mGs inhibited the somatic cAMP elevation less strongly than Lyn11-mGs.

### The cilia-localized D1R pool is preferentially ‘coupled’ to the elevation of somatic PKA activity

We next utilized the same strategy to assess the effect of targeted inhibition on the elevation of somatic PKA activity, monitored with ExRai-AKAR2 in the soma (**Fig 3A**). SKF induced a rapid and robust elevation of somatic PKA activity in the control (‘No Block’) condition (**Fig 3B**, black) that was similarly and strongly suppressed both in the ‘PM Block’ (**Fig 3B**, orange) and ‘Cilium Block’ conditions (**Fig 3B**, blue), as verified both by integrated AUC (**Fig 3C**) and peak analyses (Supplemental Fig 3A). We also observed significant suppression of the ISO-induced PKA activity elevation by both inhibitor constructs (**Fig 3D-F** and Supplemental Fig 3B).

**Figure 3.**
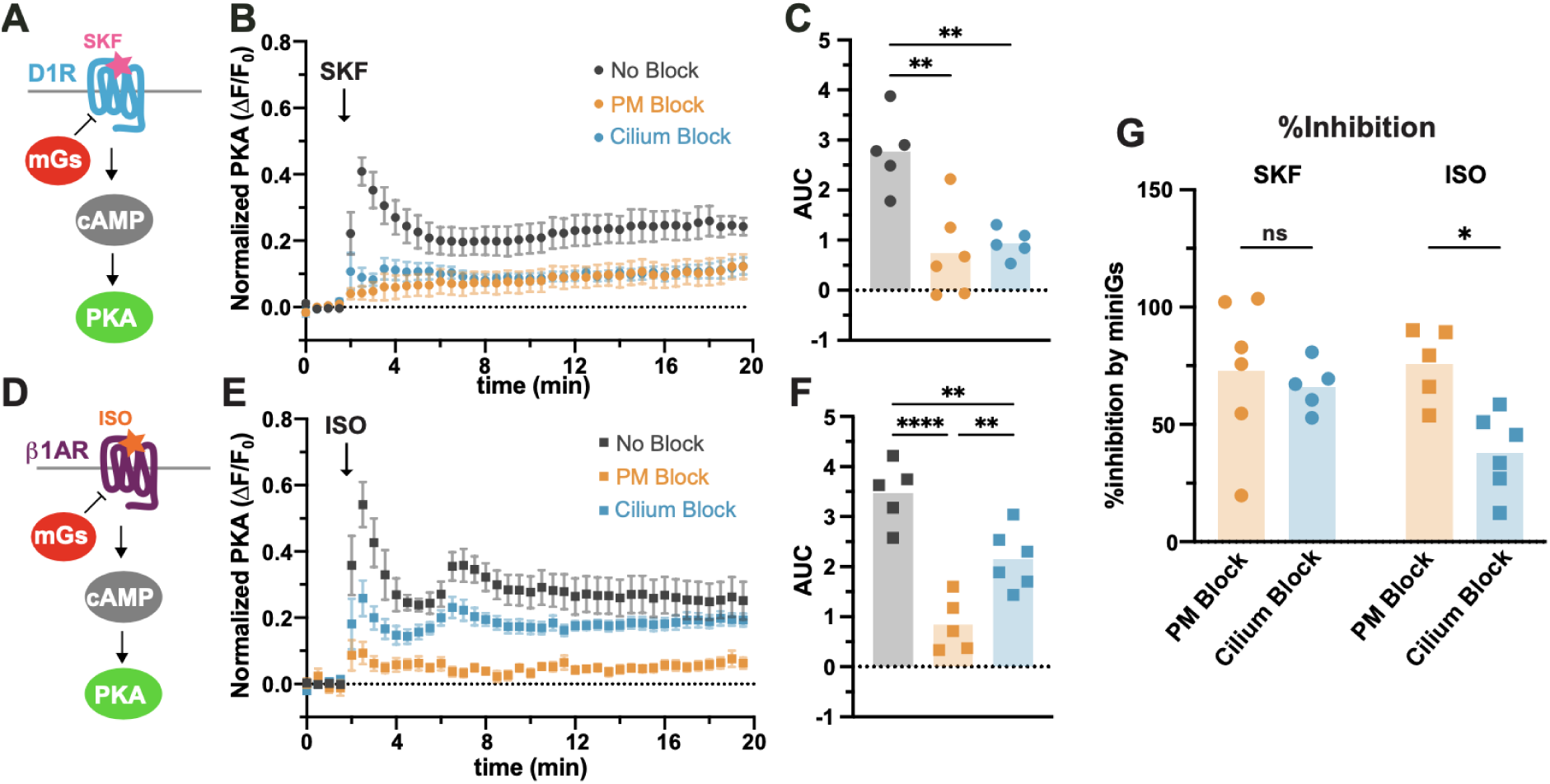
Targeted ciliary D1R blockade reveals preferential ciliary control of somatic PKA activation. **A-C.** Cultured MSNs expressing PKA biosensor (ExRai-AKAR2) and Lyn11-mApple (‘No Block’), Lyn11-mGs (‘PM Block’), or Arl13b-mGs (‘Cilium Block’) were stimulated with 1 µM SKF and fluorescence was measured and quantified as described in Figure 1. **B.** Time course of PKA response to 1 µM SKF added at 2 minutes in cultured MSNs expressing the different mGs blockers or control plasmid. **C.** Expression of the ‘PM Block’ and ‘Cilium Block’ significantly reduced the integrated PKA response to SKF in the first 5 min of agonist in MSNs. One-way ANOVA ***p = 0.0008 with Tukey’s multiple comparison test; data are presented as mean from 5-6 dishes from 6 independent neuron preps, 73-85 cells. **D-F.** Cultured MSNs expressing PKA biosensor and ‘No Block’, ‘PM Block’, or ‘Cilium Block’ were stimulated with 100 nM ISO and fluorescence was measured and quantified as described in Figure 1. **E.** Time course of PKA response to 100 nM ISO added at 2 minutes in cultured MSNs expressing the different mGs blockers or control plasmid. **F. ‘**PM Block’ and ‘Cilium Block’ both significantly reduced the integrated PKA response to ISO in the first 5 min of agonist addition in MSNs. One-way ANOVA ****p < 0.0001 with Tukey’s multiple comparison test; data are presented as mean from 5-6 dishes from 6 independent neuron preps, 54-82 cells. **G.** ‘PM’ and ‘Cilium Block’ inhibited the integrated PKA response to SKF stimulation to similar degrees, but the ‘Cilium Block’ inhibited the ISO response significantly less than the ‘PM Block’. Two-way ANOVA (interaction p = 0.10; main effect of miniGs *p = 0.02; main effect of agonist p = 0.17) with Sidak’s multiple comparisons test. *p < 0.05, **p < 0.01, ***p < 0.001, ****p < 0.0001.

While these inhibitor effects on the somatic PKA activity elevation generally paralleled results from our cAMP measurements, we noticed that the specific ‘Cilium Block’ condition suppressed the D1R-mediated PKA activity elevation particularly strongly– to nearly the same degree as the ‘PM Block’ throughout the measured time course (**Fig 3B and C**). This was not the case for the β1AR-mediated response, where the PKA activity elevation was only partially inhibited in the ‘Cilium Block’ condition yet was almost fully suppressed (like the D1R response) in the ‘PM Block’ condition (**Fig 3E, F** and Supplemental Fig 3B and C). Highlighting this difference, the calculated degree of inhibition produced by the ‘PM Block’ and ‘Cilium Block’ conditions were not significantly different for the D1R-mediated PKA response but diverged for the β1AR-mediated PKA response (**Fig 3G** and Supplemental Fig 3C). Together, these results suggest that, contrary to being restricted from initiating signaling from the primary cilium, the primary cilium of MSNs is a major subcellular location from which endogenous D1Rs initiate such signaling to drive PKA activity elevation throughout the soma.

### The primary cilium physically aligns the entire cAMP - PKA signaling axis from the cilium into the soma

Because this difference in the inhibitor effects on D1R and β1AR signaling was clear at the level of the somatic PKA activity elevation, but not at the level cAMP elevation, we hypothesized that the preferential ‘coupling’ of the PKA activity elevation elicited by the cilium-concentrating D1R might reflect a proximity effect to PKA. To test this, we examined endogenous PKA localization relative to the primary cilium. We found endogenous PKA Cα, a major PKA catalytic subunit (PKAcat) that is broadly expressed in the brain including in striatal MSNs^10,56^, to localize in a punctate distribution in the somatic cytoplasm (**Fig 4A**). This is consistent with previous evidence indicating that somatic PKA is present in concentrated stores largely outside cilia and associated with intracellular structures including Golgi membrane elements and the centrosome^10,57–59^. While we failed to detect PKA accumulation within the primary cilium of MSNs, a strong accumulation was resolved near the ciliary base that colocalized with the centrosome / ciliary basal body marker FGFR10P^60^ (FOP). When examined at higher magnification (**Fig 4A**, inset), we found each adenylyl cyclase-containing primary cilium to localize within several µm of multiple extraciliary PKA stores, including the concentrated store located within <1 µm of the FOP-marked basal body. Such close apposition between the primary cilium and extraciliary PKA stores was observed in nearly all neurons imaged (**Fig 4B**) and across different neuron preparations (Supplemental Fig 4).

**Figure 4.**
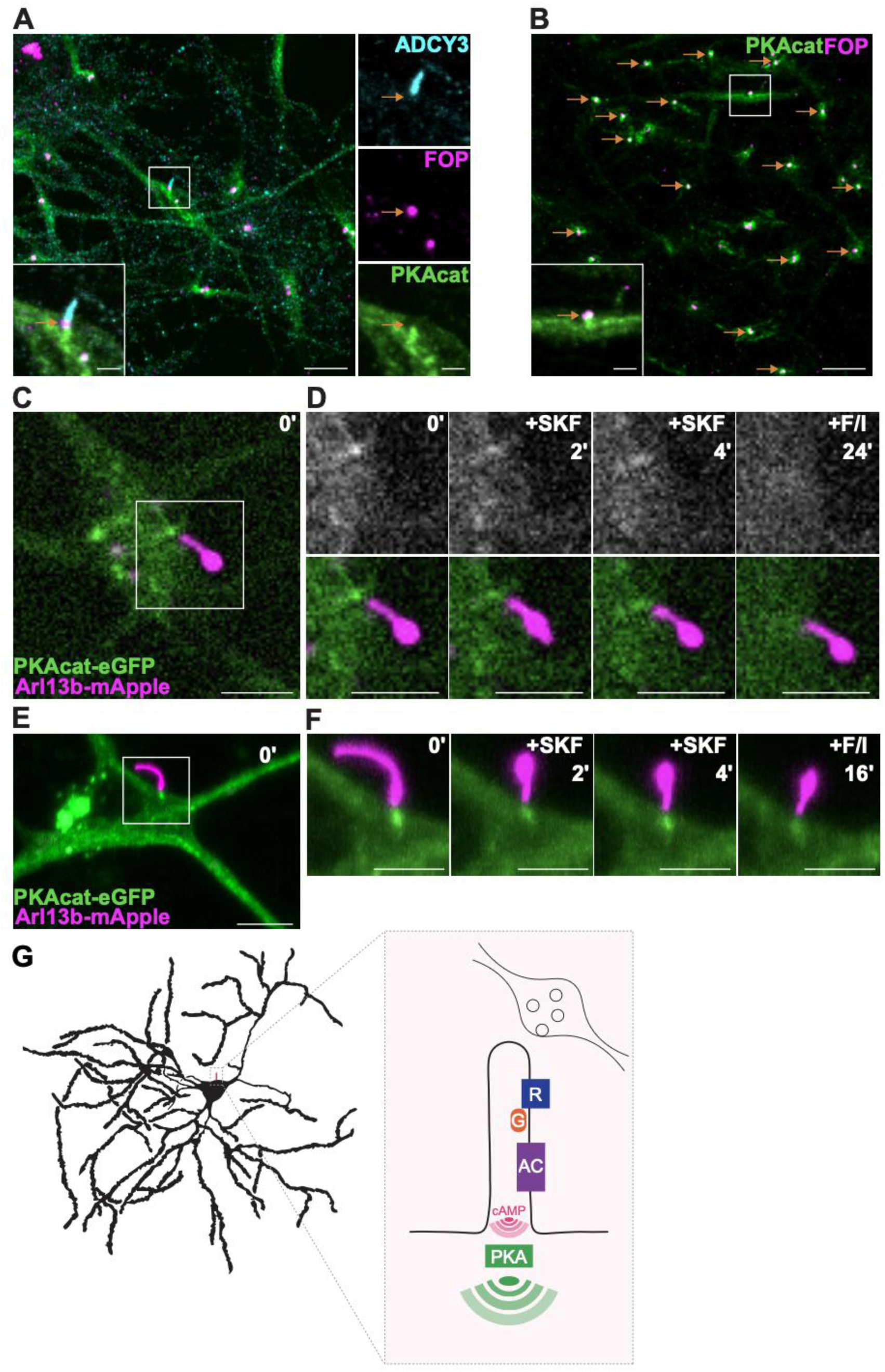
The neuronal primary cilium spatially organizes cAMP - PKA signaling into the soma. **A.** Cultured MSNs were fixed and stained for PKA catalytic domain *a* (PKAcat, green), basal body using the anti-FGFR10P antibody (FOP, magenta), and ADCY3 (cyan). Images were acquired using confocal microscopy. Arrows indicate the base of cilium with co-localized PKAcat and FOP. Scale bar: 10 µm, inset: 2 µm. **B.** Cultured striatal MSNs were fixed and stained for PKAcat (green) and FOP (magenta) to observe frequency of co-localization of PKAcat to base of cilium by confocal microscopy. See supplement for quantification. Arrows indicate cells with PKAcat in close proximity to the ciliary base. Scale bar: 10 µm, inset: 2 µm. **C.** MSNs were transfected with PKAcat-eGFP and arl13b-mApple to visualize PKAcat and primary cilia in live neurons. Image was acquired using confocal microscopy. Scale bar: 20 µm. **D.** Frames from a movie taken using a confocal microscope of the neuron in panel C treated with 1 µM SKF at 2 minutes and FSK and IBMX (F/I) at 20 minutes. Top: PKAcat-eGFP, bottom: merge with Arl13b-mApple. Scale bar: 5 µm. **E.** Cultured MSNs expressing PKAcat-eGFP and arl13b-mApple were imaged using a light sheet microscope. Scale bar: 20 µm. **F.** Frames from a movie taken using a confocal microscope of the neuron in panel **E** treated with 1 µM SKF at 2 minutes and FSK and IBMX (F/I) at 12 minutes. Top: PKAcat-eGFP, bottom: merge with arl13b-mApple. Scale bar: 5 µm. **G.** Model and schematic depicting neuromodulatory receptor signaling propagation in MSN primary cilia downstream of activation by neuromodulator released from presynaptic neuron. R: Receptor; G: G protein; AC: adenylyl cyclase.

Close proximity between the primary cilium and concentrated somatic PKA stores was also evident in living neurons, visualized using a GFP-fused version of Cα (PKAcat-eGFP) and Arl13b-mApple to mark cilia (**Fig 4C**). cAMP activates PKA by destabilizing an inhibitory interaction with regulatory subunits, resulting in the unleashing of autonomously active catalytic subunits that diffuse away from the locally regulatory subunits when cAMP rises to a sufficiently high level^25,61^. In living neurons, we found the PKAcat-eGFP accumulation resolved by confocal microscopy at the ciliary base to remain detectable throughout the time course of SKF exposure, but to dissipate after application of FSK/IBMX (**Fig 4D**). A caveat to the confocal analysis is its potential vulnerability to movement artifacts because we were able to image only a single optical plane to achieve sufficient temporal resolution without prohibitive photobleaching. To address this, we next used oblique light sheet imaging as an alternate modality that enables rapid acquisition of full cellular volumes^62^ (**Fig 4E**). This approach verified the presence of a punctate PKA accumulation near the ciliary base that persisted throughout the duration of exposure to endogenous receptor agonist, and that largely dissipated after application of FSK/IBMX (**Fig 4F**). These results suggest that the local PKA store at the ciliary base constitutes a major cAMP-responsive PKA store that can be fully mobilized under conditions of supraphysiological cAMP elevation (FSK/IBMX) but has sufficient capacity to support local activation at endogenous receptor levels.

## Discussion

By devising a method to locally inhibit GPCR activity selectively in the ciliary microdomain, the present results reveal that the primary cilium on striatal MSNs is a major site from which endogenous D1Rs initiate dopaminergic neurotransmission and propagate it through the canonical cAMP / PKA cascade into the soma. These results provide a significant advance by revising a widely held current view that the primary cilium is unable to support modulatory neurotransmission through the canonical cAMP / PKA cascade. Beyond showing that such signaling initiated by the D1R, which is known to be selectively targeted to the primary cilium, we also provide evidence suggesting that the β1AR, a close paralogue that is endogenously coexpressed in MSNs but not selectively concentrated in the ciliary microdomain, can also propagate cAMP signaling into the soma from the primary cilium. Taken together with the fact that the primary cilium comprises less than 1% of the somatic plasma membrane surface area^63^, these results suggest that the primary cilium is not only able to support modulatory neurotransmission through the canonical cAMP / PKA cascade, but that it facilitates such signaling.

Our results also suggest a simple model to explain how such transmission is achieved, using the primary cilium to propagate neurotransmission by locally concentrating key components and enforcing proximity throughout the signaling cascade– from intercalating with axonal release sites that release dopamine and initiate signaling to aligning ciliary cAMP production with major stores of PKA concentrated in the extraciliary cytoplasm (**Fig 4G**). This model of ‘long-range’ cilium-to-soma signal propagation differs from that of compartmentalized cAMP signaling thought to operate in the Hedgehog pathway, where ciliary cAMP is thought to regulate PKA activity locally in the primary cilium^29,48,64,65^, but it is fully consistent with the long-recognized presence of concentrated PKA stores in the extraciliary somatic cytoplasm, including on centrosomes^57^. We also note that our proposed model is similar to one advanced previously to explain how cAMP that is locally generated in the primary cilium directs cell migration during cortical development^66^. Here, the primary cilium and adjacent PKA stores function together as an integrated cilium-to-soma signaling unit, based on the principle of enforced spatial proximity. We propose that a similar concept is sufficient to explain how the primary cilium is able to facilitate, rather than restrict, modulatory neurotransmission through the canonical cAMP / PKA cascade in differentiated MSNs.

In addition to identifying a previously unrecognized role of the neuronal primary cilium in facilitating canonical modulatory neurotransmission through cAMP, the present results have significant implications, more broadly, for how striatal dopamine signaling is spatially organized. Dopamine has been traditionally considered to function as a diffuse ‘broadcast’ neuromodulator in the striatum, due to the high density of dopaminergic axon varicosities present there and the ability of dopamine to diffuse rapidly in brain tissue. However, accumulating evidence suggests that striatal dopamine can produce more localized effects^39^, particularly on dendrites where limited ‘hotspots’ of transient dopamine elevation have been resolved near synapses^67–69^. The primary cilium on MSNs is remarkably long, similar in length to the somatic diameter in intact striatal tissue. Based simply on the physics of diffusion^70^, this predicts that the primary cilium of MSNs is poised to considerably increase the extracellular volume that is effectively sampled by the soma. Thus, in contrast to modulatory neurotransmission onto dendrites that increasingly appears to be spatially organized, with local hotspots of release adjacent to individual synapses, the primary cilium may enable the soma to sample a much larger extracellular volume and thereby receive a distinct signal by effectively integrating the contributions of multiple axon release sites.

A key limitation of the present study, and an important future direction, is linking the presently described cell biology to specific regimes of physiological receptor activation and their outcome(s) *in vivo*. Nevertheless, considering the central importance of cAMP / PKA signaling in striatal dopaminergic neurotransmission^23^, we think it likely that the present observations will prove to have relevance. Along this line, we note that a recent study investigating the behavioral effects of genetic ablation of primary cilia in the adult mouse striatum did not reveal an overall loss of dopaminergic neuromodulation. Rather, it identified a variety of deficits among specific motor and motor learning tasks that share a dependence on the animal’s precise perception of time^71^. Because striatal dopaminergic neurotransmission is well-known to encode temporal information^39^, and in light of the present evidence indicating that the primary cilium is poised to efficiently sample dopamine and propagate its effects into the extraciliary cytoplasm, we speculate that the ‘long-range’ neurotransmission described here may contribute to temporal coordination within individual neurons or among neurons in synaptic networks.

In closing, the present results extend the concept of ciliary neurotransmission to modulatory transmission mediated by dopamine receptors in striatal MSNs. In addition they reveal that, in these neurons, the primary cilium facilitates, rather than restricts, neurotransmission through the canonical cAMP cascade.

## Methods

### Materials and reagents

pCAGGS-SSF was constructed using pGCGFP-G418 (a gift from Andrew Pierce, Addgene plasmid #31264) modified to contain a signal sequence (SS)-Flag epitope cassette resulting in a construct called ‘pCAGGS-SSF’. Arl13b fusion constructs purchased from Twist Bioscience were cloned into pGCGFP-G418 in place of the GFP coding sequence using In-Fusion (Takara) cloning. pCAGGS-Lyn11-mApple-miniGs was a gift from N. Fisher, and miniGs was removed from the plasmid using PCR to generate the pCAGGS-Lyn11-mApple construct. Flag HTR6 and ꞵ1AR plasmids were constructed using receptor cDNA purchased from Origene and inserted into pCAGGS-SSF by PC and InFusion cloning. All constructs were validated by DNA sequencing.

### Recombinant DNA

|  |  |  |
| --- | --- | --- |
| **Plasmid:** pCAGGS-SSF | This paper |  |
| **Plasmid:** Flag-β1AR | This paper | |
| **Plasmid:** Flag-D1R | Ripoll et al., |  |
| **Plasmid:** ExRai-AKAR2 | Ripoll et al., Jin Zhang |  |
| **Plasmid:** Arl13b-ExRai-AKAR2 | This paper |  |
| **Plasmid:** Lyn11-mApple | Fisher and von Zastrow |  |
| **Plasmid:** Lyn11-mApple-miniGs | This paper |  |
| **Plasmid:** Arl13b-mApple | This paper |  |
| **Plasmid:** Arl13b-mApple-miniGs | This paper |  |
| **Plasmid:** PKAcat-GFP | Ripoll et al., Peng et al., |  |
| **Plasmid:** Flag-HTR6 | This paper |  |

### Primer Table

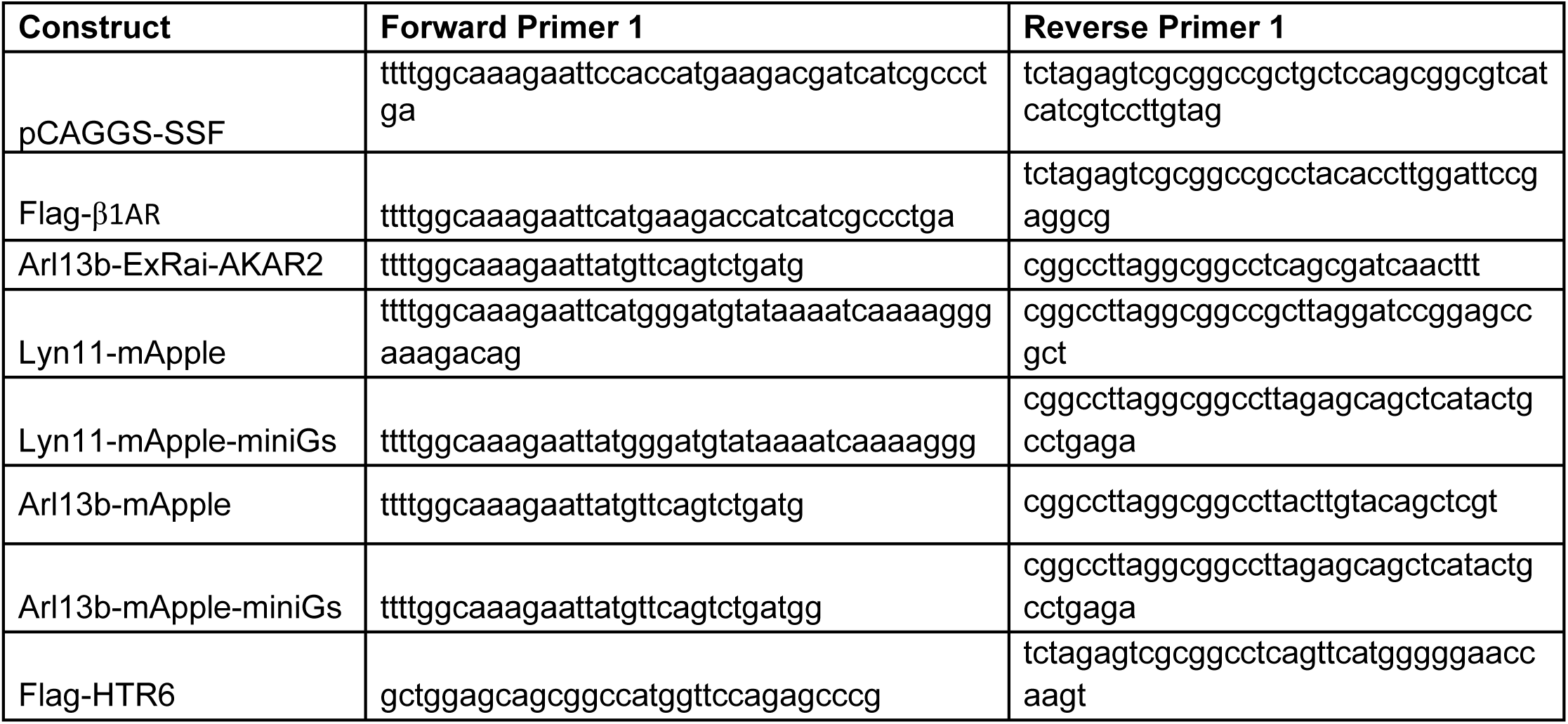

### Antibodies Table

| **Antibody** | **Purpose** | **Species** | **Dilution** | **Company** | **Product #** |
| --- | --- | --- | --- | --- | --- |
| Anti-Flag (conjugated: M1-647) | Immunostainin g | mouse | 1:1000 | Sigma-Aldrich | F-2040 |
| Anti-PKAα cat | Immunostainin g | Rabbit | 1:1000 | Santa Cruz | sc-903 |
| Anti-ARL13B | Immunostainin g | rabbit | 1:2000 | ProteinTech | 17711-1-AP |
| Anti-ACIII | Immunostainin g | rabbit | 1:1000 | Abcam | Ab125093 |
| Anti-DAT | Immunostainin g | Rat | 1:1000 | Millipore | MAB369 |
| Anti-ACIII | Immunostainin g | chicken | 1:5000 | EncorBio | CPCA_ACIII |
| Anti-ARL13B | Immunostainin<br>g | mouse | 1:1000 | ProteinTech | 66739-1-Ig |
| Anti-Flag | Immunostainin<br>g | rabbit | 1:1000 | Sigma-Aldrich | Cat#F-7425 |
| Anti-Flag ( M1) | Immunostainin<br>g | mouse | 1:1000 | Sigma-Aldrich | F-2040 |
| Anti-FGFR10P (FOP) | Immunostainin<br>g | mouse | 1:1000 | Abnova | H00011116 |

### Bacterial and virus strains

|  |  |  |
| --- | --- | --- |
| E. coli StellarTM Competent Cells | Takara | Cat#636766 |

### Chemicals, peptides, and recombinant proteins

|  |  |  |
| --- | --- | --- |
| Forskolin | Sigma-Aldrich | Cat#F6886-25mg; |
| IBMX | Sigma-Aldrich | Cat#I5879-1G |
| SKF-81297 | Tocris | Cat#1447 |
| Isoproterenol | Sigma-Aldrich | cat# I6504 |

### Critical Commercial assays

|  |  |  |
| --- | --- | --- |
| cAMP Assay: Green Up cADDIS cAMP Assay Kit (#U0200G) | Montana Molecular | https://montanamolecular.com/livecell-camp-assay-caddis/ |

### Experimental models: Organisms/strains

|  |  |  |
| --- | --- | --- |
| Rat: Sprague Dawley | Charles River | RRID:RGD_734476 |

### Software and algorithms

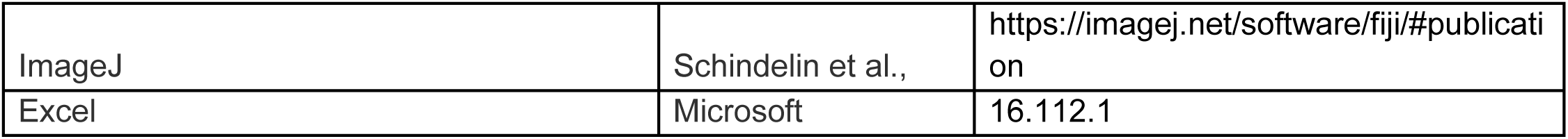

|  |  |  |
| --- | --- | --- |
| ImageJ | Schindelin et al., | https://imagej.net/software/fiji/#publicati<br>on |
| Excel | Microsoft | 16.112.1 |

### Primary neuronal culture and transfection

Primary striatal cultures were prepared from embryonic day 18 Sprague-Dawley rats (Charles River) dissected from a timed-pregnant female following euthanasia as previously described^10^. All procedures were performed according to the National Institutes of Health Guide for Care and Use of Laboratory Animals and approved by the University of California San Francisco Institutional Animal Care and Use Committee. The striatum was identified using a stereomicroscope and dissected into ice-cold Hank’s buffered saline solution calcium/magnesium/phenol red free (Thermo Fisher Scientific cat# 14-175-145). Striata were dissociated in 0.05% trypsin/EDTA (Gibco) for 15 min at 37°C and washed 2x in Dulbecco’s modified Eagle’s medium (DMEM, Gibco) supplemented with 10% fetal bovine serum (FBS, University of California, San Francisco, Cell Culture Facility) and 30 mM HEPES (Gibco). Neurons were further dissociated by trituration through flame-polished Pasteur pipettes. Neurons were seeded onto 35 mm glass bottom dishes with 20 mm micro-well (Cellvis cat# D35-20-1.5-N) coated with poly-D-lysine (Sigma). Medium was exchanged on day *in vitro* 4 (DIV4) for phenol red free Neurobasal medium (Gibco) supplemented with Glutamax 1X (Gibco) and B27 1X (Gibco). On DIV8, half of the cultured medium was exchanged with fresh, equilibrated medium. Cultures were transfected using Lipofectamine 2000 (Thermo Fisher) on DIV8 using 2 µL of Lipofectamine and 1-2 µg of DNA per 35 mm dish plated with 2x10^5^ neurons. Media was changed 4-6 hr following transfection. For PKAcat-GFP localization and signaling assays, neurons were transfected via electroporation before initial plating on DIV0. Briefly, 5-6x10^6^ neurons per condition were electroporated with up to 5 µg total DNA using Rat Amaxa Transfection kit (Lonza, cat# VPG-1003), and 2-4x10^5^ neurons were plated onto 35 mm glass bottom dishes. All neurons were cultured in a humidified incubator with 5% CO_2_ at 37°C throughout, and imaged between DIV11 and DIV15. All experiments were performed with at least 3 independent cultures.

### Immunostaining of Fixed Cultured Neurons

To label surface Flag-tagged receptors expressed in striatal neurons, on DIV12 neurons were incubated with mouse M1 anti-flag antibody (see key resources table) diluted 1:1000 in media, 15 min, 37°C. Neurons were then washed with phosphate-buffered saline with calcium and magnesium (PBS, Life Technologies cat# 14040216) and fixed for 10 min at RT in 4% paraformaldehyde (PFA) and 4% sucrose /PBS (Fisher Scientific cat# AAJ19943K2). Neurons were blocked in 0.2% Triton X-100 and 3% normal goat serum (CST, cat# 5425S) diluted in PBS for 1 hr at RT. Primary antibodies were diluted in blocking solution and incubated overnight at 4°C rocking gently. Neurons were then washed three times in 0.2% Triton X-100 in PBS, 10 min, at RT. Secondary antibodies were diluted in blocking solution and labeling was performed at RT for 1 hr. Cells were washed one time, 10 min with 0.2% Triton X-100 in PBS, labeled for 10 min with DAPI (Sigma)/PBS (diluted 1:20,000), and stored in 1X PBS. Neurons were imaged by confocal microscopy using a Nikon Ti inverted fluorescence microscope with CSU-22 spinning disk confocal with a 60 × 1.4 numerical aperture oil objective.

### Immunostaining of Fixed Mouse Brain Sections

Mice were euthanized by carbon dioxide inhalation and transcardially perfused with PBS followed by 4% PFA/PBS (ThermoFisher) as detailed previously^37^. 30% sucrose in PBS was used as cryoprotectant. Cryoprotected brains were embedded in O.C.T. (Sakura Finetek) and sectioned at 30 μm using a Cryostat (Leica CM3050 S). Sections were rehydrated in PBS prior to blocking in 3% normal goat serum, 0.3% Triton X-100 diluted in PBS at RT for 1 hr. Primary antibodies were incubated overnight followed by 3 washes with PBS-T. Secondary antibodies were added for 2 hr at RT (See key resources table). Slices were mounted using ProLong Diamond (ThermoFisher Scientific), imaged using a Nikon 6D/High Throughput inverted fluorescence microscope with CSU-W1 large field view with a 100 × 1.4 numerical aperture oil objective, and stitched using automated image stitching 10x.

### Confocal fluorescence microscopy

For live neuron imaging, media was exchanged with Hepes Buffered Solution (HBS, 120 mM NaCl, 2 mM KCl, 10 mM HEPES, 2 mM CaCl_2_, 2 mM MgCl_2_, 0.9 g D-glucose, pH 7.4) prior to bringing the dish to the microscope. Neurons were imaged on an inverted Nikon Ti2-E spinning disk confocal microscope equipped with a CREST X-Light V2 large field of view camera using a 40x 1.4 NA oil objective. Neurons were kept at 37°C in a temperature- and humidity-controlled chamber.

### Light sheet fluorescence microscopy

Cultured neurons were maintained under live-cell imaging conditions and imaged using a custom-built ‘Snouty’ single objective light sheet microscope constructed by T. Fadero, equipped with a Hammamatsu Orca-Quest-2 camera and a 60x 1.3 NA Si objective. Excitation was achieved using a 405/488/561/639 nm single mode fiber laser launch (Vortran VersaLase). Emission was collected using single-pass filter sets (Chroma) verified to spectrally separate fluorescent channels with negligible bleedthrough under the imaging conditions used. Neurons were imaged on coverslips maintained in HBS imaging medium at 37°C in a humidified 5% CO₂ environmental chamber (Oko). Full-volume cell images were acquired every 20 seconds under the control of MicroManager^72^.

### Neuronal signaling assays using fluorescent biosensors

Live cell cAMP assay was conducted using cADDis Green Up cAMP biosensor (Montana Molecular) and live cell PKA activity assay was conducted using ExRai-AKAR2^44^ using single 488nm laser illumination as described previously.^10^ Endogenous D1R or β1AR were selectively activated by bath application of 1 μM SKF 81297 hydrobromide (SKF, Tocris cat# 1447) or 100 nM isoproterenol hydrochloride (ISO, Sigma Aldrich, cat# I6504), respectively, after a 2 min baseline. Images were acquired every 20 - 30 seconds as indicated. 10 μM forskolin (Fsk, Millipore Sigma) combined with 500 µM 3-isobutyl-1-methylxanthine (IBMX, Millipore Sigma) was added at the end of each imaging episode (typically at 18 min) and acquisition was continued for an additional 5 min to record the maximal (saturated) biosensor response for that episode. Image analysis was conducted using FIJI (ImageJ) on unprocessed images by integrating the fluorescent signal measured in ROIs manually drawn around individual cells or cilia, as indicated, and subtracting background fluorescence defined by that measured in an adjacent area not containing a fluorescent neuron. Normalized cAMP (cADDis) and PKA activity (ExRai-AKAR2) responses were determined by dividing the background-subtracted fluorescence by the maximum background-subtracted fluorescence signal determined after Fsk/IBMX addition. Integrated cAMP and PKA responses were measured as the area under the curve of two phases (0-5 and 0-18 min of agonist treatment) when indicated. The peak value corresponds to the maximum fluorescence value between 0 and 5 min after SKF or ISO addition.

### DAT varicosities and bouton quantification

The number of DA release sites in close proximity to striatal cilia were estimated by counting the number of varicosities labeled with the DAT antibody within approximately 1 micron of an ADCY3-labeled cilium in the mouse striatal brain slices. This was done for 20 cilia from one z-projection confocal image using ImageJ Cell Counter Macro.

### Cilia localization and enrichment quantification

Quantitative image analysis was performed on unprocessed 16-bit files using FIJI software (ImageJ). Receptor-positive cilia were counted in at least 10 images per condition and receptor(+) cilia were determined based on whether a cell expressed the receptor and was ciliated. Unciliated cells were not counted. To measure cilia enrichment, a region of interest (ROI) was defined using thresholding on the cilia marker channel. This ROI was applied to the receptor channel to determine relative signal of the receptor within the cilium and then moved to the plasma membrane region to measure receptor expression on the cell surface. The ROI was moved to a region outside of the cell to correct for background signal and this value was subtracted from both cilia and plasma membrane values. Relative cilia enrichment was determined by dividing the corrected cilia signal by plasma membrane signal.

### Statistical analysis and reproducibility

Results are displayed as a mean of at least 3 independent experiments ±SEM and include multiple samples per experiment. Statistically significant differences were determined using One-way or Two-way ANOVA (for more than two comparisons) with appropriate multiple comparisons test (see Figure Legends), Pearson correlation or unpaired t-test was performed using Prism 9 software (GraphPad Software). Statistical significance is denoted as: ns (not significant, p > 0.05), *p < 0.05, **p < 0.01, ***p < 0.001, ****p < 0.0001.

## Data availability

Source data are provided with this paper, and additional data and materials are available upon reasonable request from the corresponding authors.

## Acknowledgements

We thank Max Nachury, Jeremy Reiter, Markus Delling, Adraham Beyene, José Velilla and Mia Konjkusic for valuable discussion, David Julius for access to the cryomicrotome, Nicole Fisher for critical comments on the manuscript and all members of the Ehrlich and von Zastrow groups for useful discussion and support. We thank Tanner Fadero for building the oblique light sheet microscope and providing instruction and advice on its operation, with financial support from the UCSF Program for Breakthrough Biomedical Research (PBBR), and members of the UCSF Center for Advanced Light Microscopy (Kari Herrington and Julia Martin) for valuable advice, instruction and assistance. Much of the imaging data for this study were acquired at the UCSF Center for Advanced Light Microscopy directed by Nico Stuurman. RRF received support from an NIH Ruth Kirchstein Postdoctoral Fellowship (F32MH130096), XJH from an NIH/NRSA BRAIN Initiative Postdoctoral Fellowship (F32MH136662) and ATE from an NIH Mentored Research Career Development Award (K01MH123757). ATE also received support from the Brain and Behavior Research Foundation (award #27303). These studies were supported by research grants from the NIH (R01MH120212, R01DA010711 and R01DA012864 to M.v.Z) and a collaborative research award from the UCSF Program in Breakthrough Biomedical Research.

## Author contributions

RRF, ATE and MvZ conceptualized the study and designed experiments. XJH contributed to the design of signaling experiments and the data analysis pipeline. NS oversaw experiments using the light sheet microscope, and designed and implemented the image rendering pipeline. RRF, ATE and XJH performed experiments and analyzed data. RRF, ATE and MvZ wrote the manuscript with input from all authors.

## Supplemental Figures

**Supplemental Figure 1:**
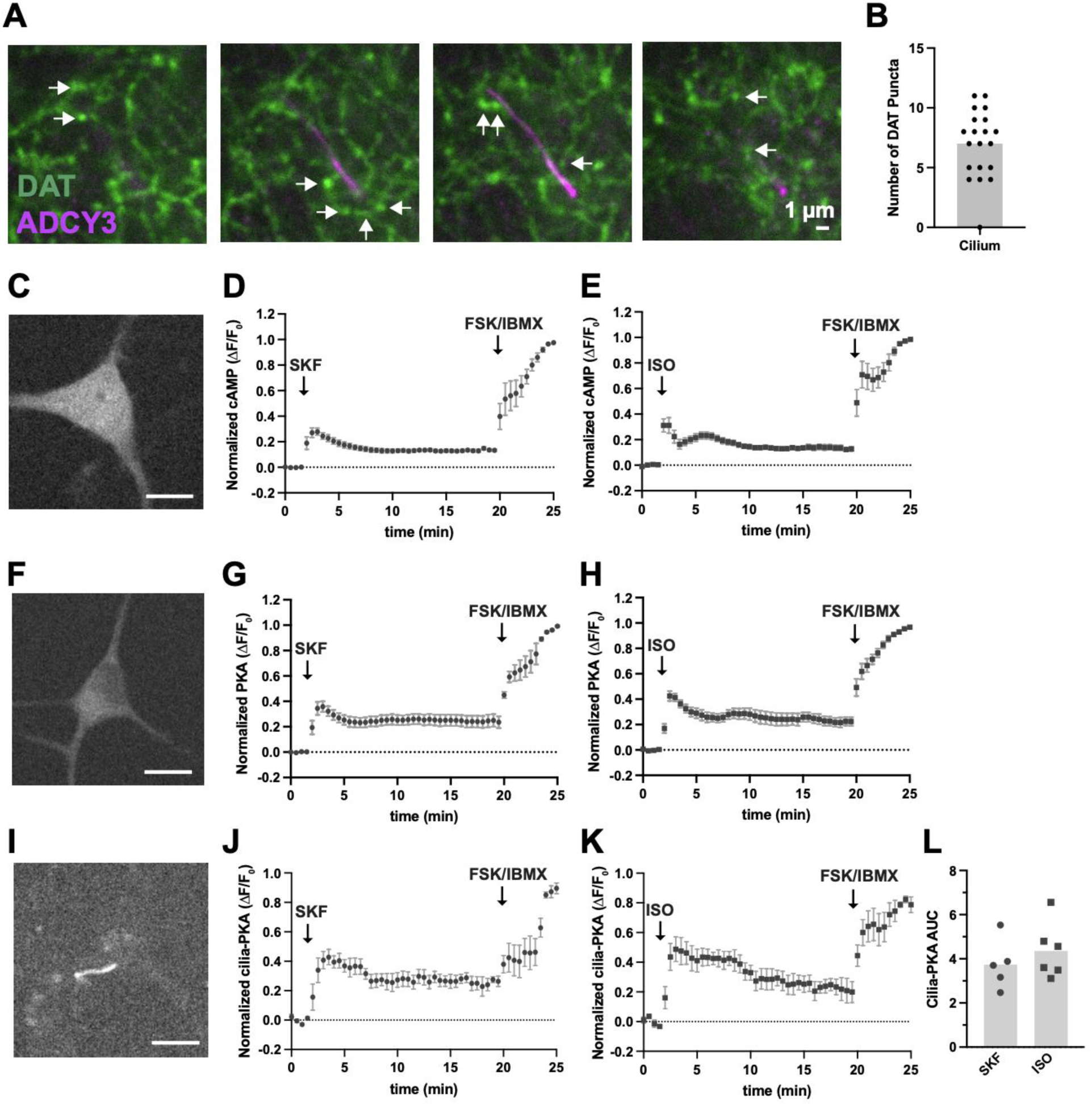
Multiple dopamine terminals are within 1 µm of striatal neuron primary cilia and validation of cAMP and PKA biosensors for measuring cAMP - PKA signaling by native receptors in striatal MSNs. **A.** Higher magnification of the highlighted region from Fig 1A depicting how DAT puncta were counted at different z-planes of the original image (neuronal primary cilia (ADCY3), dopamine terminals (DAT), and nuclei (DAPI) in adult mouse dorsal striatum). Arrows indicate DAT puncta. Scale bar: 1 µm. **B.** Average number of DAT puncta within 1 micron of 20 ADCY3-labeled primary cilia from Fig 1A. **C.** Representative micrograph of cultured MSN expressing cAMP biosensor, cADDis. Scale bar: 10 µm. **D.** Full time course of cAMP response to 1 µM SKF from Figure 1E, including 10 µM FSK and 500 µM IBMX addition at 20 minutes. Data in Figure 1E are normalized to maximum FSK/IBMX response. **E.** Full time course of cAMP response to 100 nM ISO from Figure 1F, including 10 µM FSK and 500 µM IBMX addition at 20 minutes. Data in Figure 1F are normalized to maximum FSK/IBMX response. **F.** Representative micrograph of cultured MSN expressing PKA biosensor, ExRai-AKAR2. Scale bar: 10 µm. **G.** Full time course of PKA response to 1 µM SKF from Figure 1I, including 10 µM FSK and 500 µM IBMX addition at 20 minutes. Data in Figure 1I are normalized to maximum FSK/IBMX response. **H.** Full time course of PKA response to 100 nM ISO from Figure 1J, including 10 µM FSK and 500 µM IBMX addition at 20 minutes. Data in Figure 1J are normalized to maximum FSK/IBMX response. **I.** Representative micrograph of cultured MSN expressing the cilia PKA biosensor, cilia-ExRai-AKAR2. Scale bar: 10 µm. **J.** Full time course of cilia PKA response to 1 µM SKF at 2 minutes and 10 µM FSK and 500 µM IBMX addition at 20 minutes. **K.** Full time course of cilia PKA response to 100 nM ISO at 2 minutes and 10 µM FSK and 500 µM IBMX addition at 20 minutes. **L.** Average integrated PKA response to SKF or ISO within the first 5 minutes of agonist addition by endogenous receptors in cultured MSNs. Data are presented as mean from 5-6 dishes from 4-5 independent neuron preps, 41-50 cells.

**Supplemental Figure 2:**
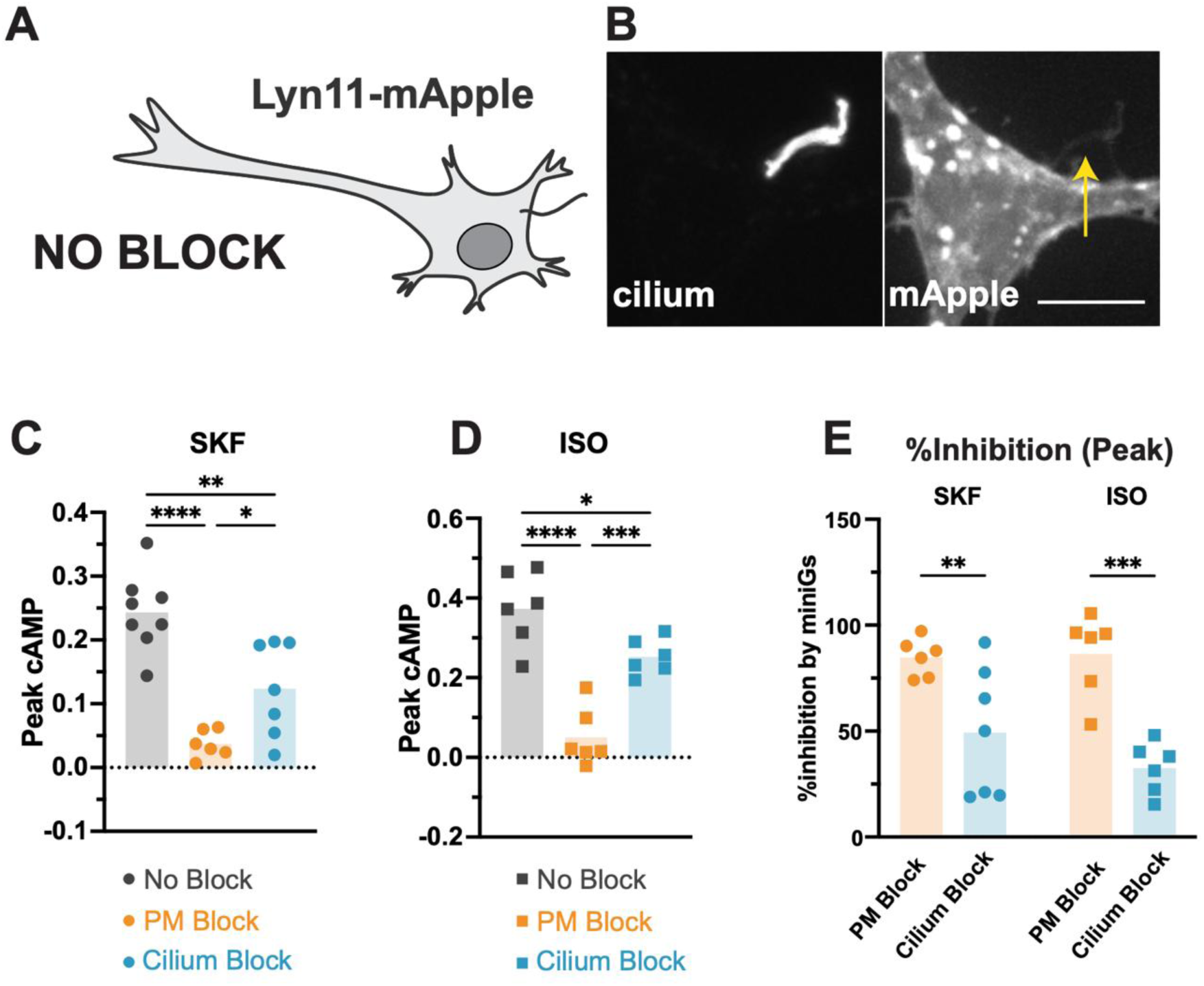
Extended analysis of compartment-specific receptor blockade of receptor-evoked cAMP responses. **A.** Schematic describing the localization of the Lyn11-mApple ‘No Block’ control construct in neurons. **B.** Representative images of Lyn11-mApple co-expressed with Flag-HTR6 (left) to identify cilia. lyn11-mApple is expressed along the plasma membrane, including the ciliary membrane. Arrow indicates mApple localized to primary cilia. Scale bars: 10 µm. **C.** Peak cAMP response within the first 5 minutes of SKF treatment from the time course in Figure 2D. ‘PM Block’ and ‘Cilium Block’ significantly reduced the peak response to SKF in MSNs, and peak response to SKF was inhibited significantly more by ‘PM Block’ than ‘Cilium Block’. One-way ANOVA ****p < 0.0001 with Tukey’s multiple comparison test; data are presented as mean from 6-8 dishes from 7 independent neuron preps, 65-170 cells. **D.** Peak response within the first 5 minutes of ISO treatment from the time course in Figure 2G. ‘PM Block’ and ‘Cilium Block’ significantly reduced the peak response to ISO in MSNs, and peak response to ISO was inhibited significantly more by ‘PM Block’ than ‘Cilium Block’. One-way ANOVA ****p < 0.0001 with Tukey’s multiple comparison test; data are presented as mean from 6 dishes from 6-7 independent neuron preps, 65-103 cells. **E.** The % inhibition of the peak cAMP response was compared between SKF and ISO treatments. The ‘Cilium Block’ inhibited peak cAMP significantly less than ‘PM Block’ for both SKF and ISO. Two-way ANOVA (interaction *p = 0.27; main effect of mGs ****p < 0.0001; main effect of agonist p = 0.36) with Sidak’s multiple comparisons test. *p < 0.05, **p < 0.01, ***p < 0.001, ****p < 0.0001.

**Supplemental Figure 3:**
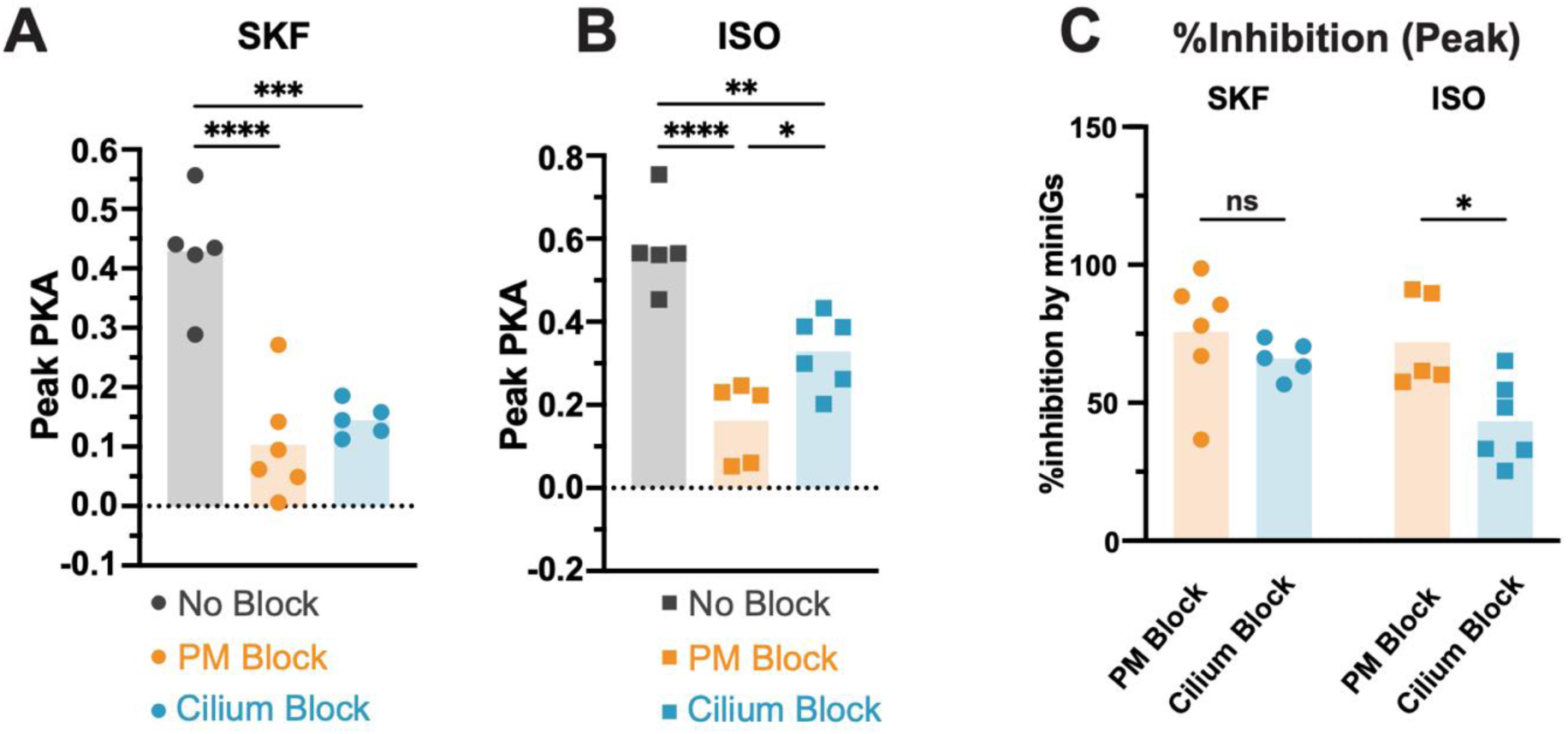
Extended analysis of PKA responses following compartment-specific receptor blockade. **A.** Peak PKA response within the first 5 minutes of SKF treatment from the time course in Figure 3B. Expression of the PM Blocker and Cilium Blocker significantly reduced the peak response within the first 5 min in MSNs. One-way ANOVA Peak ****p < 0.0001 with Tukey’s multiple comparison test; data are presented as mean from 5-6 dishes from 6 independent neuron preps, 73-85 cells. **B.** Peak PKA response within the first 5 minutes of ISO treatment from the time course in Figure 3E. Expression of the PM Blocker and Cilium Blocker significantly reduced the peak response within the first 5 min in MSNs. One-way ANOVA Peak ****p < 0.0001 with Tukey’s multiple comparison test; data are presented as mean from 5-6 dishes from 6 independent neuron preps, 54-82 cells. **C.** The % inhibition of the peak PKA response was compared between SKF and ISO treatments. ‘Cilium Block’ and’ PM Block’ inhibited the peak PKA response to SKF similar degrees, but the ‘Cilium Block’ inhibited the ISO response significantly less than the ‘PM Block’. Two-way ANOVA (interaction *p = 0.19; main effect of miniGs *p = 0.01; main effect of agonist p = 0.08) with Sidak’s multiple comparisons test. *p < 0.05, **p < 0.01, ***p < 0.001, ****p < 0.0001.

**Supplemental Figure 4:**
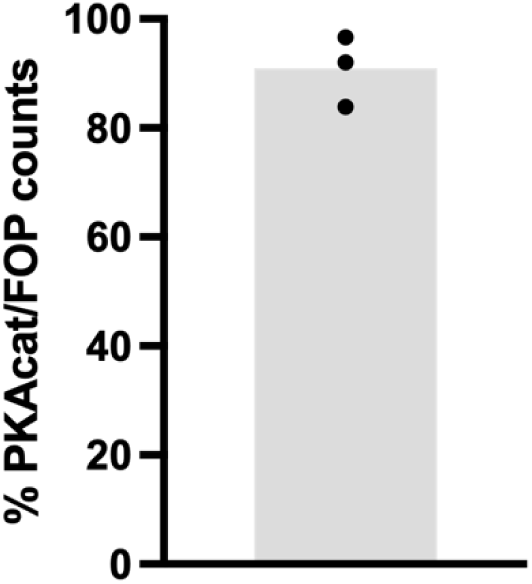
PKAcat co-localizes with basal bodies in striatal MSNs. Quantification of immunostained striatal neurons stained with anti-PKAcat to label concentrated PKA stores and anti-FGFR10P (FOP) to label basal bodies. The percentage of MSNs with co-localized PKAcat puncta and FOP basal bodies relative to the total number of MSNs containing FOP basal bodies is shown for n = 3 independent experiments, with 56–108 cells analyzed per experiment.

